# Rapid cold hardening modifies mechanisms of ion regulation to delay anoxia-induced spreading depolarization in the CNS of *Locusta migratoria*

**DOI:** 10.1101/2023.04.27.538621

**Authors:** Phinyaphat Srithiphaphirom, Yuyang Wang, Maria J. Aristizabal, R. Meldrum Robertson

**Affiliations:** Department of Biology, Queen’s University, Kingston, ON, K7L 3N6, Canada

**Keywords:** Locust, Anoxia, Spreading depolarization, Rapid cold hardening, Potassium transport

## Abstract

Insects live in varied habitats and experience different kinds of environmental stresses. These stresses can impair neural performance, leading to spreading depolarization (SD) of nerve cells and neural shutdown underlying coma. The sensitivity of an insect’s nervous system to stress (e.g., anoxia) can be modulated by acute pre-treatment. Rapid cold hardening (RCH) is a form of preconditioning, in which a brief exposure to low temperature can enhance the stress tolerance of insects. SD is associated with a sudden loss of ion, notably K^+^, homeostasis. We used a pharmacological approach to investigate whether RCH affects anoxia-induced SD in the locust, *L. migratoria*, via one or more of the following homeostatic mechanisms: (1) Na^+^/K^+^-ATPase (NKA), (2) Na^+^/K^+^/2Cl^-^ co-transporter (NKCC), and (3) voltage-gated K^+^ (K_v_) channels. We also assessed abundance and phosphorylation of NKCC using immunoblotting. We found that inhibition of NKA or K_v_ channels delayed the onset of anoxia-induced SD in both control and RCH preparations. However, NKCC inhibition preferentially abrogated the effect of RCH. Additionally, we observed a higher abundance of NKCC in RCH preps but no statistical difference in its phosphorylation level, indicating the involvement of NKCC expression or degradation as part of the RCH mechanism.

## 1. Introduction

Insects are found in all land and freshwater habitats, thus they are exposed to and challenged by natural environmental stresses (e.g., anoxia, extreme temperatures) and likely experience drastic changes in body temperature, hydration state, and oxygen availability, which present significant physiological challenges. To survive, insects have adapted mechanisms of tolerance that allow them to overcome these challenges. The ability of insects to tolerate periods of stress is highly dependent on the sensitivity of their nervous system to unfavourable conditions. This sensitivity is plastic and can be modulated by acute pre-treatments (e.g., brief exposure to environmental stressors such as high and low temperature, anoxia, and dehydration) (Coulson and Bale, 1991; Gantz et al., 2020; Hou et al., 2014; Lee et al., 1987; Rodgers et al., 2007; Srithiphaphirom et al., 2019; Srithiphaphirom and Robertson, 2022).

Rapid cold hardening (RCH) is a form of thermal acclimation in which brief chilling (minutes to hours) significantly enhances the stress tolerance of an animal (see *review* (Teets et al., 2020)). RCH occurs in many invertebrate species and has been studied intensively in insects (e.g., (Armstrong et al., 2012; Czajka and Lee, 1990; Gantz et al., 2020; Lee et al., 1987; Owen et al., 2013; Srithiphaphirom et al., 2019; Srithiphaphirom and Robertson, 2022)). In flesh flies (*Sarcophaga crassipalpis* and *Sarcophaga bullata*), ̴ 90% of larvae and adults do not survive at −10 °C, however, pre-exposure to 0 °C for 2 hours increases their survival to 91.1% at −10 °C (Lee et al., 1987). RCH at 4 °C for 4 hours delays chill coma and enhances anoxic tolerance by increasing the time to succumb to an anoxic environment (water submersion) in intact locusts, *L. migratoria* (Gantz et al., 2020; Srithiphaphirom et al., 2019). Proposed mechanisms underlying RCH include accumulation of cryoprotectants (e.g., glycerol, sorbitol), modification of cell membrane, and changes in gene and protein expression (e.g., (Chen et al., 1987; Michaud and Denlinger, 2006, 2007; Overgaard et al., 2005)). Although the benefits of RCH in improving insect performance are well-established, the underlying mechanisms that enhance stress-induced tolerances remain unsettled.

Due to their small size and varied habitats, insects can naturally be exposed to anoxia by suffocation during heavy rains and flooding. Anoxia disrupts the energy supply and depletion of energy leads to reduced activity of ATP-dependent ion pumps, causing dissipation of ion gradients responsible for proper neural function. To thrive, many insects enter a reversible coma, or hypo-energetic state (Campbell et al., 2018; Campbell et al., 2019; Rodgers et al., 2007), when they are under stress such as anoxia. Coma onset is driven by spreading depolarization (SD), a phenomenon characterized by a sudden loss of ion, notably potassium ion (K^+^), homeostasis, and temporary shutdown of the central nervous system (CNS) (Andersen and Overgaard, 2019; Dreier and Reiffurth, 2015; Herreras and Makarova, 2020; Leao, 1944; Robertson et al., 2020; Robertson et al., 2017; Rodgers et al., 2010; Rodgers et al., 2007). Insect comas are reversible and may provide an adaptive advantage for coping with environmental stressors. CNS arrest could conserve limited energy resources or reallocate them to more important cellular and tissue operations (Robertson et al., 2020; Robertson et al., 2023). Additionally, shutting down neural activity could protect the nervous system from further damage during stress. In insects, SD can be measured across a blood-brain barrier (BBB) formed from perineurial and subperineurial glial cells connected by septate and gap junctions (Limmer et al., 2014; Robertson et al., 2020; Schofield and Treherne, 1984; Stork et al., 2008). The insect BBB regulates ion diffusion between the insect haemolymph and the CNS interstitium to protect the underlying neurons from large changes in extracellular ion composition (Schofield and Treherne, 1984; Treherne and Schofield, 1979, 1981). Maintenance of ion homeostasis in the CNS is crucial for proper neural function.

Our recent studies showed that RCH at 4 °C for 4 hours delays SD induced by chilling (Srithiphaphirom et al., 2019) and anoxia (Srithiphaphirom and Robertson, 2022). We believe that an imbalance in processes of K^+^ accumulation and K^+^ clearance in the CNS triggers a rapid surge of extracellular K^+^ concentration in the CNS ([K^+^]_o(CNS)_), which indicates SD. Movement of ions, including K^+^, across the BBB requires transport mechanisms because the ganglion sheath has low ion permeability and the septate junctions in the sheath limit paracellular diffusion. In this study, we used a pharmacological approach to investigate whether RCH affects anoxia-induced SD in the locust, *L. migratoria*, via one or more of the following homeostatic mechanisms that are involved in maintaining K^+^ homeostasis: (1) Na^+^/K^+^-ATPase (NKA), (2) Na^+^/K^+^/2Cl^-^ co-transporter (NKCC), and (3) voltage-gated K^+^ (K_v_) channels. NKA requires ATP to pump three sodium ions (Na^+^) out of the cell in exchange for two K^+^ into the cell. For neural function, NKCC normally transports Na^+^, K^+^, and two Cl^-^ into the cell, and K_v_ channels normally allow flow of K^+^ out of the cell in response to changes in the transmembrane potential. Thus, NKA and NKCC aid K^+^ clearance, whereas K_v_ channels promote K^+^ accumulation in the extracellular space.

We used SD to indicate the onset of anoxia-induced neural shutdown that underlies coma and measured the DC potential across the BBB. First, we used ouabain to inhibit NKA, previously shown to increase K^+^ efflux across the cockroach BBB, which reflects increases in [K^+^]_o(CNS)_ (Kocmarek and O’Donnell, 2011). We found that ouabain hastened anoxia-induced SD in both control and RCH locusts and delayed recovery from SD in control locusts. Next, we used bumetanide to block NKCC. In locusts, bumetanide disrupts the regulation of [K^+^]_o(CNS)_ by impairing K^+^ clearance (Spong et al., 2015). We found that NKCC inhibition hastened anoxia-induced SD only in RCH locusts. Moreover, immunoblotting revealed an increase in the abundance of NKCC following RCH but no difference in protein phosphorylation level. Finally, we blocked K_v_ channels using tetraethylammonium (TEA), a general blocker of K_v_ channels, or 4-aminopyridine (4-AP), specific for *Shaker* K_v_ channels. In locusts, the [K^+^]_o(CNS)_ surge during heat-induced SD is partially via TEA-sensitive K^+^ channels (Rodgers et al., 2007) and the *Shaker* family of K_v_ channels is associated with cold acclimation (Bayley et al., 2020). We found that TEA delayed anoxia-induced SD in control locusts and had an additive effect on RCH, whereas 4-AP affected neither control nor RCH locusts. Moreover, TEA was the only pharmacological agent in this study that affected the amplitude of DC potential. These findings lead us to conclude that in the locust CNS, NKA and K_v_ channels (excluding the *Shaker* family) modulate SD occurrence and possibly take part in the mechanism of RCH, whereas NKCC is directly involved in the mechanism of RCH.

## 2. Methods

### 2.1. Animals

*L. migratoria* were raised in a crowded colony maintained in the Department of Biology at Queen’s University. Locusts were reared in large, well-ventilated cages under a 12 hour:12 hour light:dark cycle with the temperature inside the cages at 25 ± 1 °C (light) and 21 ± 1 °C (dark) and fed daily with fresh wheat seedlings and a dry mixture of skim milk powder, torula yeast, and wheat bran. We randomly selected adult locusts, 4 – 5 weeks past imaginal ecdysis from the colony. We performed all experiments on both male and female locusts in a 1:1 ratio.

### 2.2. RCH pre-treatment

Treatment parameters were chosen to be consistent with our previous studies on RCH in locusts (Srithiphaphirom et al., 2019; Srithiphaphirom and Robertson, 2022). RCH pre-treatment consisted of 2 steps: (1) RCH locusts were held in ventilated plastic containers (̴ 750 mL; 1 locust per container) and maintained at approximately 4 °C for 4 hours in a cold room and (2) RCH locusts were then maintained at room temperature (RT; 22 ± 2 °C) for 1 hour before the experiment to allow locust body temperature to return to RT for recovery. Control locusts were held in similar containers (̴ 750 mL; 1 locust per container) at RT for 5 hours before the experiment.

### 2.3. Electrophysiology

#### 2.3.1. Dissected preparation

We used semi-intact preparations for measuring electrical activity as previously described (Robertson and Pearson, 1982). The legs, wings, and pronotum were removed and a dorsal midline incision was made. The locust was pinned dorsal side up onto a corkboard and the gut, air sacs, and fat bodies were removed to expose the thoracic ganglia. No nerve roots were cut.

We added standard locust saline containing (in mM): 147 NaCl, 10 KCl, 4 CaCl_2_, 3 NaOH, and 10 HEPES buffer (pH 7.2) (all chemicals from Sigma-Aldrich https://www.sigmaaldrich.com) to the thoracic and abdominal cavities. A chlorided silver wire was placed in the posterior tip of the abdomen to ground the preparation (**Figure 1A**).

**Figure 1.**
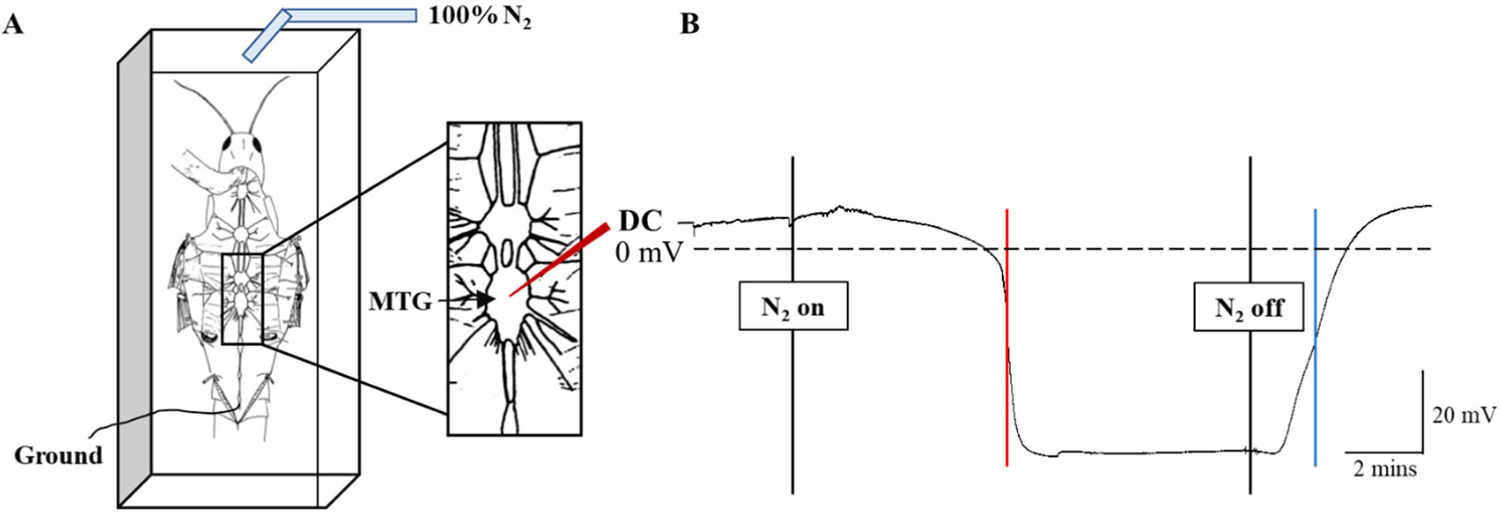
Semi-intact preparation of the locust during anoxia. **A**: a schematic diagram of semi-intact locust with recording microelectrode placement in the metathoracic ganglion (MTG) to measure DC potential (red in inset). **B**: sample extracellular recording of DC potential in the MTG relative to saline ground during exposure to anoxia using nitrogen gas (N_2_). A dotted line in **B** indicates 0 mV. A red line and a blue line in **B** indicate the point where the DC potential went negative at half-maximal amplitude and the point where the DC potential returned to the baseline level at half-maximal amplitude, respectively.

#### 2.3.2. Electrophysiological recordings

SD is characterized by an abrupt development of a negative DC potential that is coincident with the surge in extracellular K^+^ concentration in the CNS ([K^+^]_o(CNS)_) (Rodgers et al., 2007). We measured DC potential extracellularly from a single location within the metathoracic ganglion (MTG) using microelectrodes prepared from 1 mm diameter, filamented borosilicate glass capillary tubes (World Precision Instruments; WPI, Sarasota, FL, USA) pulled to form low resistance tips (approximately 5–7 MΩ) using a model P-87 Flaming/Brown micropipette puller (Sutter Instruments, Novato, CA, USA). These microelectrodes were backfilled with 500 mM KCl. Before inserting through the sheath of the MTG, we adjusted the microelectrode recording to 0 mV in the saline bathing the preparation. The signals were amplified and digitized using a model 1600 Neuroprobe amplifier (A-M Systems Inc., Sequim, WA, USA) and a Digidata 1550A (100 kHz sampling rate; Molecular Devices Inc., Sunnyvale, CA, USA), and displayed and saved using AxoScope 10.7 (Molecular Devices Inc.) (**Figure 1B**).

#### 2.3.3. Anoxia

Anoxia was used to induce neural shutdown. A nitrogen gas (N_2_) supply was placed at the end of an acrylic chamber (5 x 2.5 x 2 cm). We placed the locust pinned onto the corkboard in the chamber and partially sealed the chamber with electrical tape. Room air and N_2_ travelled through a modified glass pipette hooked over the end of the chamber. Prior to anoxia, an aquarium pump pumped room air through the chamber at ∼100 mL/min. We switched from room air to 100% N_2_ from a pressurized cylinder at ∼1 L/min to induce anoxia. The locust was exposed to N_2_ until the negative DC potential occurred rapidly, indicating the onset of anoxia-induced SD. N_2_ flow was maintained for 5 minutes after the rapid negative DC potential occurred. Then we switched back to room air to allow recovery and the recovery period was recorded for 5 minutes.

#### 2.3.4. Pharmacology

Collagenase, ouabain, bumetanide, tetraethylammonium (TEA) chloride, and 4-aminopyridine (4-AP) were obtained from Sigma-Aldrich. We used standard locust saline to prepare all pharmacological agents. Collagenase was used in this study to soften the neural lamella (Schofield et al., 1984) and aid penetration of electrodes and pharmacological agents into the ganglia. 200 µL of 1% collagenase was added to the semi-intact locust and left for 10 minutes. Then saline was used to wash out collagenase before adding any pharmacological agents. The concentrations of ouabain, TEA, and 4-AP and treatment duration of ouabain were decided based on dose-response curves (**Supplementary Figures S1 – S3**). The concentration of bumetanide and treatment durations of bumetanide, TEA, and 4-AP were used as previously described (Rodgers et al., 2007; Spong et al., 2015).

### Ouabain

To inhibit NKA, 200 µL of 5×10^-3^ M ouabain was bath-applied to the thoracic ganglia for 10 minutes before exposing the preparation to anoxia. Ouabain can induce SD in insects (e.g., (Rodgers et al., 2009; Spong et al., 2015)). However, this concentration and treatment duration of ouabain did not induce SD within the time course of the experiment.

### Bumetanide

To block NKCC, 200 µL of 10^-3^ M bumetanide was bath-applied to the thoracic ganglia for 20 minutes before exposing the preparation to anoxia.

### Tetraethylammonium

TEA is a widely used blocker of K_v_ channels. 200 µL of 5×10^-2^ M TEA was bath-applied to the thoracic ganglia for 15 minutes before exposing the preparation to anoxia.

### 4-aminopyridine

4-AP is a relatively selective blocker of members of the *Shaker* family of K_v_ channels. 200 µL of 5×10^-4^ M 4-AP was bath-applied to the thoracic ganglia for 15 minutes before exposing the preparation to anoxia.

### 2.4. Immunoblotting

#### 2.4.1. Dissection

The locusts were restrained, and the limbs were removed. We made a ventral incision between the meso- and meta-sternum to create a small opening, and the MTG was extracted and stored in 2 mL microcentrifuge tubes. Individual locust dissection and tissue collection were completed within 90 seconds and each ganglion was snap-frozen in liquid nitrogen. All tissue samples were stored at – 80 °C.

#### 2.4.2. Protein extraction

For each sample, MTGs from 4 locusts were pooled. The total protein was isolated using the TRIzol reagent (Invitrogen, Waltham, MA, USA) following the manufacturer’s instructions. Ganglia in 1 mL of TRIzol reagent were homogenized using a TissueLyzer II bead mill (one 5 mm stainless steel bead per tube, 30 s^-1^ frequency for 3 minutes; Qiagen Inc. Hilden, Germany). The homogenates were incubated with 200 µL chloroform and centrifuged for 15 minutes at 12000 x g to isolate the protein-containing organic phase. The total protein pellet was obtained by adding 1.5 mL of isopropanol to the organic phase and centrifuging for 10 minutes at 12000 xg. The pellet was washed twice with 0.3 M guanidine hydrochloride in 95% ethanol and once in 100% ethanol, before being resuspended in 170 µL of 1% SDS. The samples were treated with a phosphatase inhibitor cocktail (10 mM NaPPi, 5 mM EDTA, 5 mM EGTA, 5 mM NaF, 0.1 mM sodium orthovanadate) and cOmplete ULTRA protease inhibitor cocktail (Sigma-Aldrich) and incubated at 50 °C for 30 minutes to aid pellet resuspension. The protein samples were then stored at −20 °C for no more than a week. The total protein content in samples was obtained using a BCA protein assay kit (Sigma-Aldrich) with the SpectraMax Paradigm microplate reader (Molecular Devices, San Jose, CA, USA).

#### 2.4.3. SDS-PAGE

To determine the optimal protein amount for immunoblotting, a standard curve was generated and analyzed using the Empiria 2.0 software linear range detection function (LI-COR Biosciences, Lincoln, NE, USA). 20ug of protein were resolved on a 7.5% SDS-PAGE gel and transferred to onto a nitrocellulose membrane (0.45uM pore size). Total protein signal was obtained using the Revert 700 total protein stain kit (LI-COR) prior to evaluating signal for the C-terminus of NKCC1/2 (MABS1237, 0.5 µg/mL) or N-terminal phosphorylation site of NKCC1 (ABS1004, 1:1000). Membranes were scanned on an Odyssey XF imaging system and signal was quantified using the Empiria 2.0 software (LI-COR).

### 2.5. Statistical analysis

We analyzed the electrical recordings using Clampfit 10.7 (Molecular Devices Inc.). For each recording, we determined the time to SD, the recovery time from SD, and the amplitude of the DC potential shift. The time to SD refers to the time taken from when N_2_ was turned on to the half-maximal amplitude of the negative shift of DC potential. The recovery time from SD refers to the time taken from when N_2_ was turned off to the half-maximal amplitude of the DC potential when returning to the positive baseline level. The amplitude of the DC potential shift refers to the difference between the stable baseline level and the 2.5-minute point after the DC potential dropped. Then, we used SigmaPlot 12.5 (Systat Software Inc., Chicago, IL, USA) to visualize data and conduct statistical analyses. First, we tested data for normality using the Shapiro-Wilk test and equal variance using the Brown-Forsythe test. We used Three-way ANOVAs to identify statistical significance within and among groups (sex, treatment, and drug) for the electrophysiology experiments and Two-way ANOVAs to identify statistical significance within and between sex and treatment groups for the immunoblotting experiments. After the Three-way ANOVA and the Two-way ANOVA revealed no effect of sex, the results are presented with pooled data from both sexes. All-pairwise post hoc analysis was used to determine the significance between treatments within each measure (Holm-Sidak). We report data as the means and standard errors (SE) for parametric data or as the medians and interquartile ranges (IQR) for non-parametric data (**Supplementary Tables S1 & S2**). P < 0.05 is indicated by symbols on the figures. Asterisk (⁎) indicates a statistically significant difference within treatment and dagger (†) indicates a statistically significant difference within drug. The plots are overlaid with individual data points plotted as open symbols.

## 3. Results

### 3.1. Ouabain hastens anoxia-induced SD in both control and RCH locusts and delays recovery from SD in control locusts

In the ouabain experiments, the results of time to SD did not fit the assumptions of a parametric test, thus we transformed the data by taking log_10_x. There was no effect of sex (Three-way ANOVA: p = 0.649), no interaction between treatment and sex (Three-way ANOVA: p = 0.635), and no interaction between drug and sex (Three-way ANOVA: p = 0.334) in both control and RCH pre-treatments. In control pre-treatment, the time to SD of locusts with ouabain was significantly shorter than the time to SD of locusts with saline (Holm-Sidak: p < 0.001) (**Figure 2A**). Similarly, in RCH pre-treatment, the time to SD of locusts with ouabain was significantly shorter than the time to SD of locusts with saline (Holm-Sidak: p < 0.001) (**Figure 2A**). Within drugs, the time to SD of RCH locusts with saline was significantly longer than the time to SD of control locusts with saline (Holm-Sidak: p < 0.001) (**Figure 2A**). The time to SD of RCH locusts with ouabain was also significantly longer than the time to SD of control locusts with ouabain (Holm-Sidak: p = 0.008) (**Figure 2A**).

**Figure 2.**
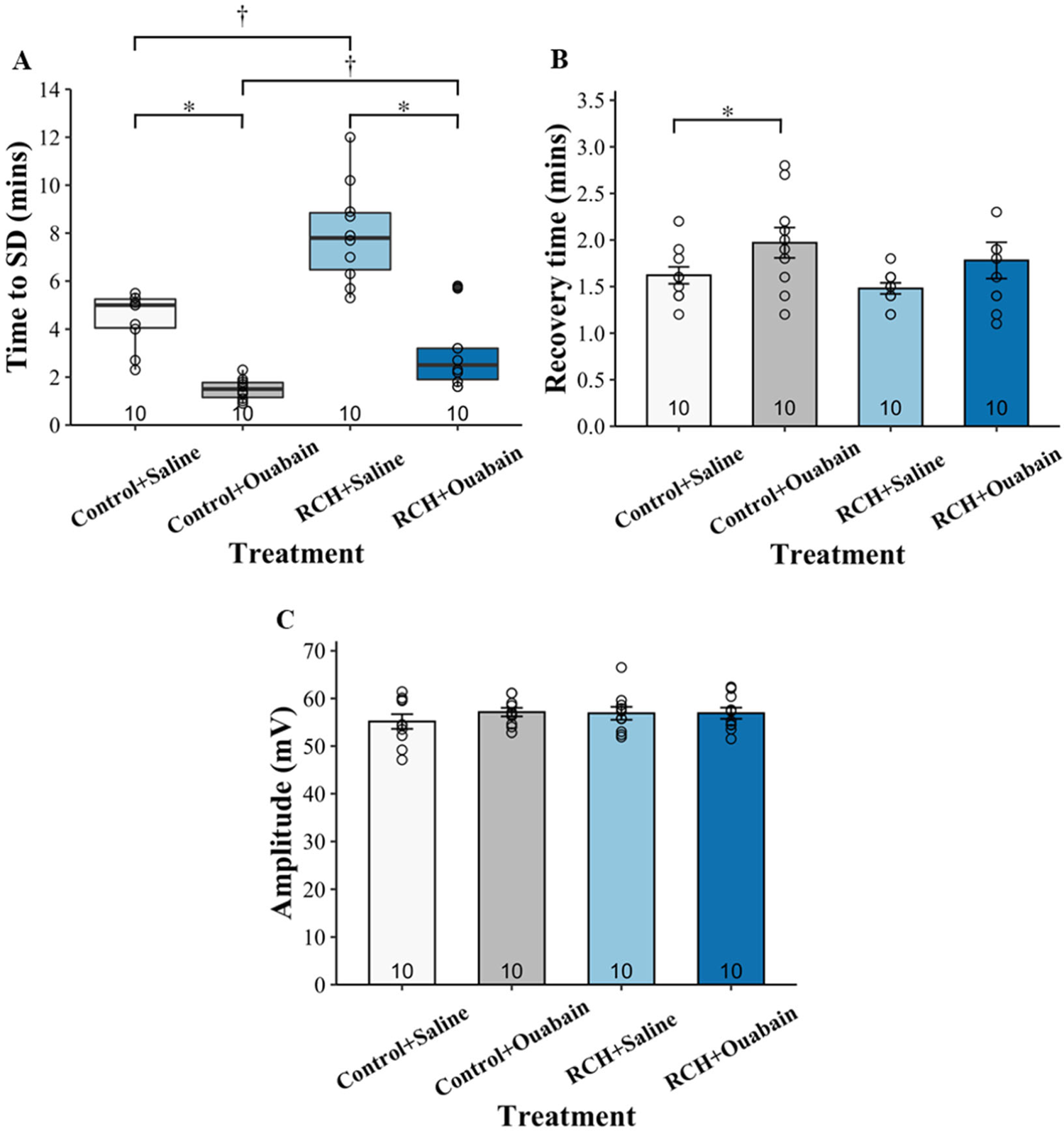
The effect of ouabain. **A**: ouabain hastens anoxia-induced SD by reducing time to SD of both control and RCH locusts (statistical tests performed with transformed data, see text). **B, C**: ouabain increases recovery time from SD of control locusts but does not affect amplitude of DC potential of control and RCH locusts. Data are medians, IQR, and means ± S.E. (n = 10 each group; p-values are reported in Results). For graphical display, data from the two sexes are pooled as no sex differences were observed. Asterisk (⁎) indicates a statistically significant difference within treatment (i.e., Control or RCH). Dagger (†) indicates a statistically significant difference within drug (i.e., Saline or Ouabain).

For the recovery time from SD and the amplitude of DC potential, there was no effect of sex (Three-way ANOVA: recovery time p = 0.801; amplitude p = 0.887), no interaction between treatment and sex (Three-way ANOVA: recovery time p = 0.128; amplitude p = 0.946), and no interaction between drug and sex (Three-way ANOVA: recovery time p = 0.746; amplitude p = 0.570) in both control and RCH pre-treatments. Ouabain increased the recovery time from SD of control locusts (Holm-Sidak: p = 0.038) but did not significantly change the recovery time from SD of RCH locusts (Holm-Sidak: p = 0.159) (**Figure 2B**). There was no significant difference in amplitudes of DC potential of control locusts with saline, control locusts with ouabain, RCH locusts with saline, and RCH locusts with ouabain (Three-way ANOVA: p = 0.465; **Figure 2C**).

### 3.2. Bumetanide hastens anoxia-induced SD only in RCH locusts

In the bumetanide experiments, there was no effect of sex (Three-way ANOVA: p = 0.303), no interaction between treatment and sex (Three-way ANOVA: p = 0.202), and no interaction between drug and sex (Three-way ANOVA: p = 0.535) in both control and RCH pre-treatments. In control pre-treatment, the times to SD of locusts with saline and locusts with bumetanide were not significantly different (Holm-Sidak: p = 0.547) (**Figure 3A**). However, in RCH pre-treatment, the time to SD of locusts with bumetanide was significantly shorter than the time to SD of locusts with saline (Holm-Sidak: p = 0.001) (**Figure 3A**). Within drugs, the time to SD of RCH locusts with saline was significantly longer than the time to SD of control locusts with saline (Holm-Sidak: p < 0.001) (**Figure 3A**). The time to SD of RCH locusts with bumetanide was also significantly longer than the time to SD of control locusts with bumetanide (Holm-Sidak: p = 0.003) (**Figure 3A**).

**Figure 3.**
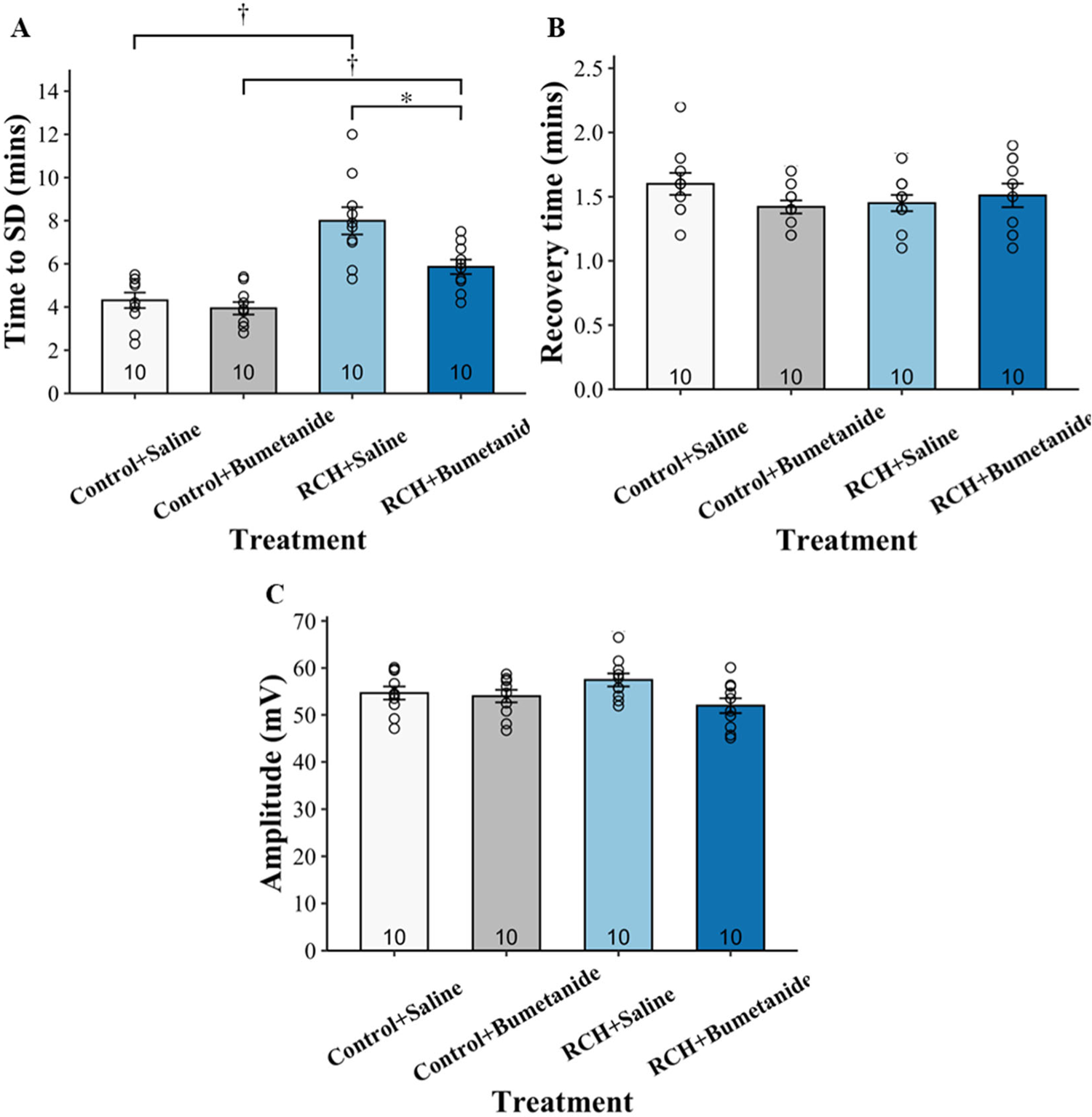
The effect of bumetanide. **A**: bumetanide hastens anoxia-induced SD by reducing time to SD of RCH locusts but does not affect time to SD of control locusts. **B, C:** bumetanide does not affect recovery time from SD and amplitude of DC potential of control and RCH locusts. Data are means ± S.E. (n = 10 each group; p-values are reported in Results). For graphical display, data from the two sexes are pooled as no sex differences were observed. Asterisk (⁎) indicates a statistically significant difference within treatment (i.e., Control or RCH). Dagger (†) indicates a statistically significant difference within drug (i.e., Saline or Bumetanide).

For the recovery time from SD and the amplitude of DC potential, there was no effect of sex (Three-way ANOVA: recovery time p = 0.791; amplitude p = 0.990), no interaction between treatment and sex (Three-way ANOVA: recovery time p = 0.791; amplitude p = 0.793), and no interaction between drug and sex (Three-way ANOVA: recovery time p = 0.063; amplitude p = 0.614) in both control and RCH pre-treatments. There was no significant difference in recovery times from SD (Three-way ANOVA: p = 0.118; **Figure 3B**) and amplitudes of DC potential (Three-way ANOVA: p = 0.119; **Figure 3C**) of control locusts with saline, control locusts with bumetanide, RCH locusts with saline, and RCH locusts with bumetanide.

### 3.3. TEA has an additive effect on RCH and reduces the amplitude of negative DC shift

In the TEA experiments, there was no effect of sex (Three-way ANOVA: p =0.406), no interaction between treatment and sex (Three-way ANOVA: p = 0.820), and no interaction between drug and sex (Three-way ANOVA: p = 0.581) in both control and RCH pre-treatments. In control pre-treatment, the time to SD of locusts with TEA was significantly longer than the time to SD of locusts with saline (Holm-Sidak: p = 0.003) (**Figure 4A**). Similarly, in RCH pre-treatment, the time to SD of locusts with TEA was significantly longer than the time to SD of locusts with saline (Holm-Sidak: p < 0.001) and was the longest among all four groups (**Figure 4A**). Within drugs, the time to SD of RCH locusts with saline was significantly longer than the time to SD of control locusts with saline (Holm-Sidak: p < 0.001) (**Figure 4A**). The time to SD of RCH locusts with TEA was also significantly longer than the time to SD of control locusts with TEA (Holm-Sidak: p < 0.001) (**Figure 4A**).

**Figure 4.**
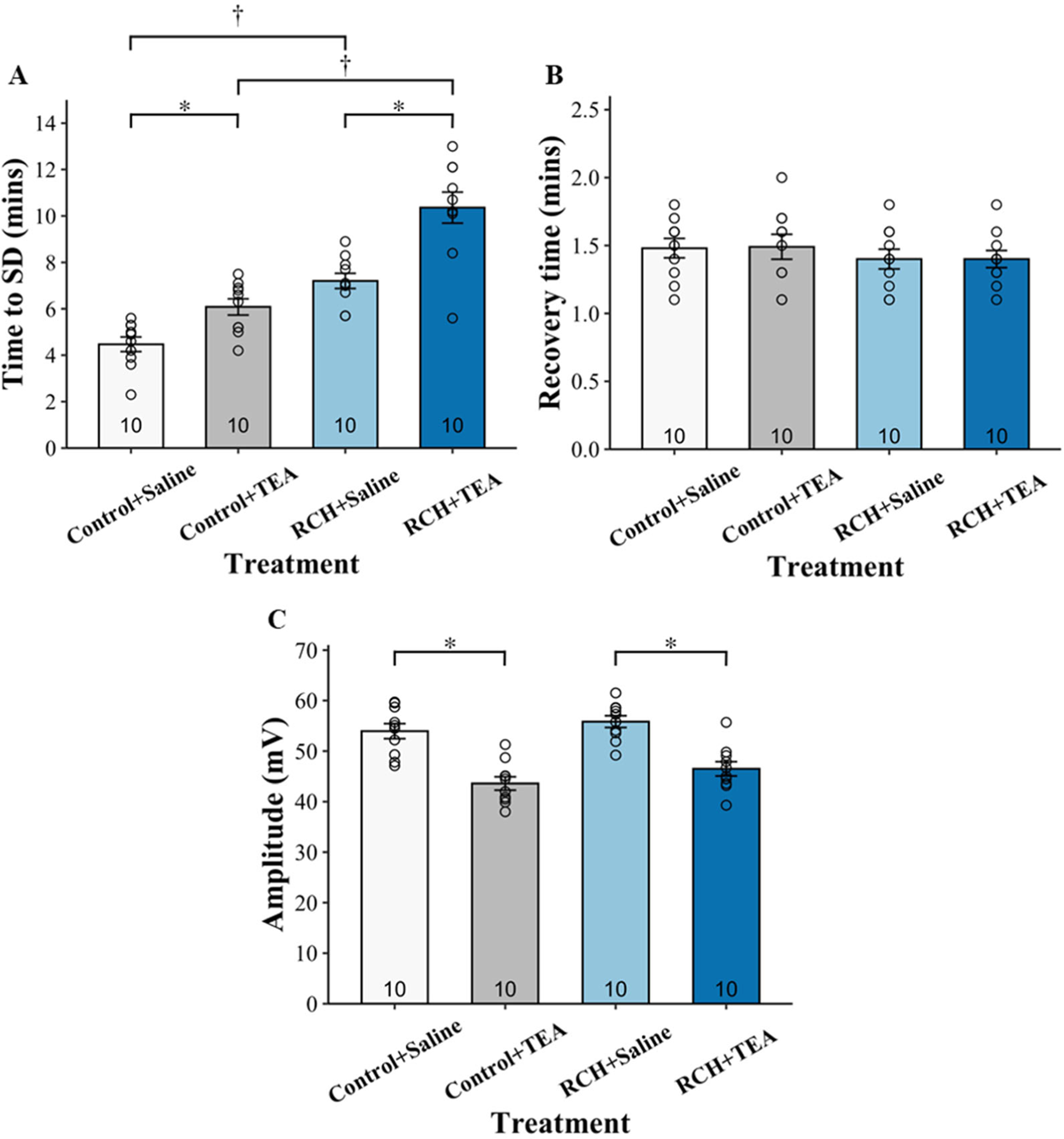
The effect of TEA. **A:** TEA delays anoxia-induced SD by increasing time to SD of control locusts and shows an additive effect on RCH locusts. **B, C:** TEA does not affect recovery time from SD but reduces amplitude of DC potential of both control and RCH locusts. Data are means ± S.E. (n = 10 each group; p-values are reported in Results). For graphical display, data from the two sexes are pooled as no sex differences were observed. Asterisk (⁎) indicates a statistically significant difference within treatment (i.e., Control or RCH). Dagger (†) indicates a statistically significant difference within drug (i.e., Saline or TEA).

For the recovery time from SD and the amplitude of DC potential, there was no effect of sex (Three-way ANOVA: recovery time p = 0.656; amplitude p = 0.663), no interaction between treatment and sex (Three-way ANOVA: recovery time p = 0.567; amplitude p = 0.243), and no interaction between drug and sex (Three-way ANOVA: recovery time p = 0.567; amplitude p = 0.912) in both control and RCH pre-treatments. There was no significant difference in recovery times from SD of control locusts with saline, control locusts with TEA, RCH locusts with saline, and RCH locusts with TEA (Three-way ANOVA: p = 0.949; **Figure 4B**). However, TEA decreased the amplitudes of DC potential of both control and RCH locusts (Holm-Sidak: p < 0.001; **Figure 4C**).

### 3.4. 4-AP affects neither control nor RCH locusts

In the 4-AP experiments, there was no effect of sex (Three-way ANOVA: p = 0.062), no interaction between treatment and sex (Three-way ANOVA: p = 0.658), and no interaction between drug and sex (Three-way ANOVA: p = 0.150) in both control and RCH pre-treatments. In control pre-treatment, the times to SD of locusts with saline and locusts with 4-AP were not significantly different (Holm-Sidak: p = 0.759) (**Figure 5A**). Similarly, in RCH pre-treatment, the times to SD of locusts with saline and locusts with 4-AP were not significantly different (Holm-Sidak: p = 0.063) (**Figure 5A**). Within drugs, the times to SD of control locusts (both with saline and with 4-AP) were significantly shorter than the times to SD of RCH locusts (both with saline and with 4-AP) (Holm-Sidak: p < 0.001) (**Figure 5A**).

**Figure 5.**
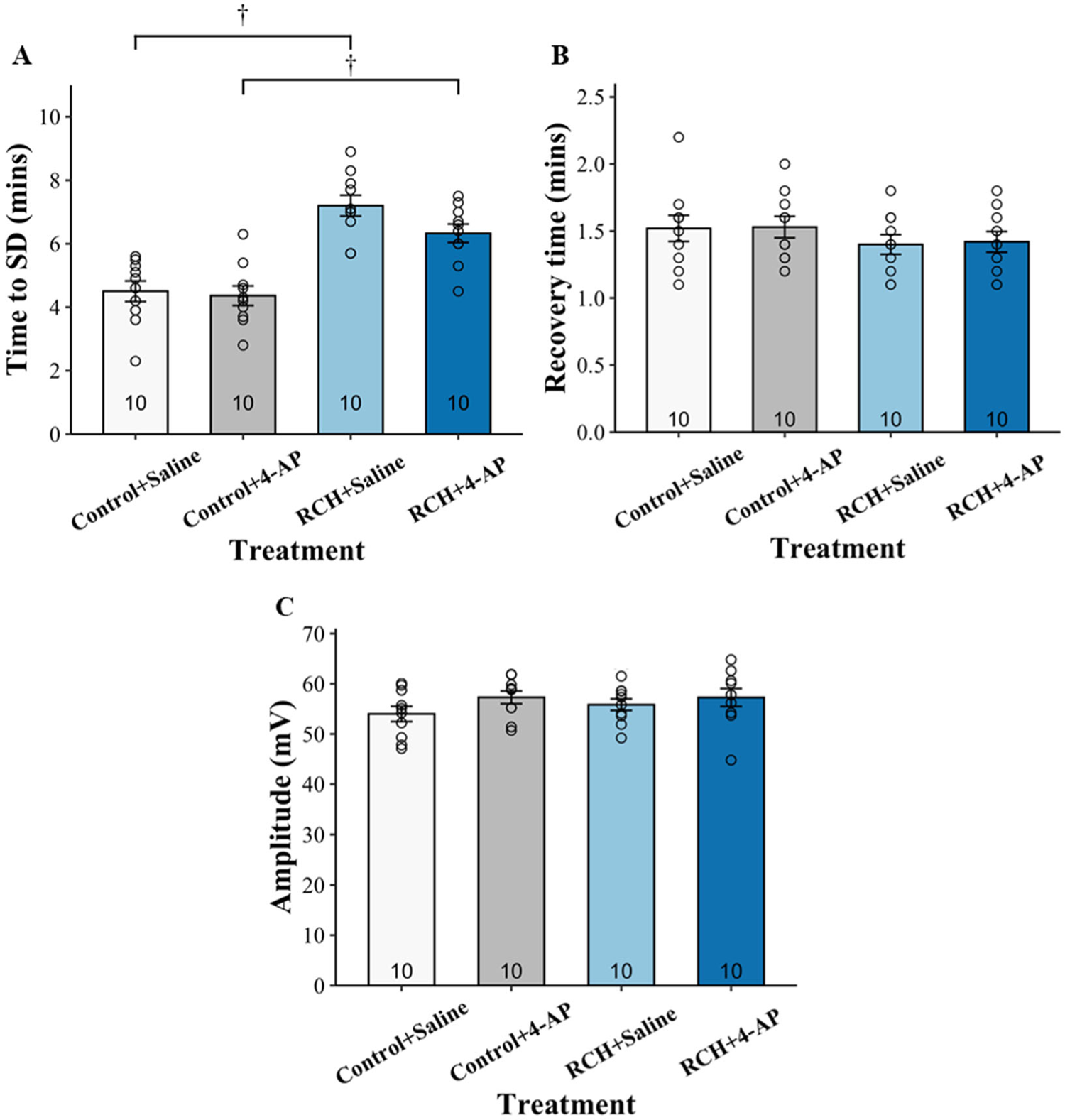
The effect of 4-AP. 4-AP does not affect **A**: time to SD, **B**: recovery time from SD, and **C**: amplitude of DC potential of both control and RCH locusts. Data are means ± S.E. (n = 10 each group; p-values are reported in Results). For graphical display, data from the two sexes are pooled as no sex differences were observed. Dagger (†) indicates a statistically significant difference within drug (i.e., Saline or 4-AP).

For the recovery time from SD and the amplitude of DC potential, there was no effect of sex (Three-way ANOVA: recovery time p = 0.515; amplitude p = 0.653), no interaction between treatment and sex (Three-way ANOVA: recovery time p = 0.678; amplitude p = 0.518), and no interaction between drug and sex (Three-way ANOVA: recovery time p = 0.443; amplitude p = 0.489) in both control and RCH pre-treatments. There was no significant difference in recovery times from SD (Three-way ANOVA: p = 0.953; **Figure 5B**) and amplitudes of DC potential (Three-way ANOVA: p = 0.535; **Figure 5C**) of control locusts with saline, control locusts with 4-AP, RCH locusts with saline, and RCH locusts with 4-AP.

### 3.5 RCH increases protein abundance of NKCC

The immunoblotting experiment detected an increased abundance of NKCC protein level in the RCH locusts (Two-way ANOVA: p = 0.006; **Figure 6A, B**); there was no effect of sex (Two-way ANOVA: p = 0.18) and no interaction between sex and treatment (Two-way ANOVA: p = 0.10). RCH had limited effect on the abundance of phospho-NKCC (pNKCC; Two-way ANOVA: p = 0.15; **Figure 6C**), there was no effect of sex (Two-way ANOVA: p = 0.29) and no interaction between sex and treatment (Two-way ANOVA: p = 0.95). Notably, a trend was observed that RCH potentially decreased NKCC phosphorylation.

**Figure 6.**
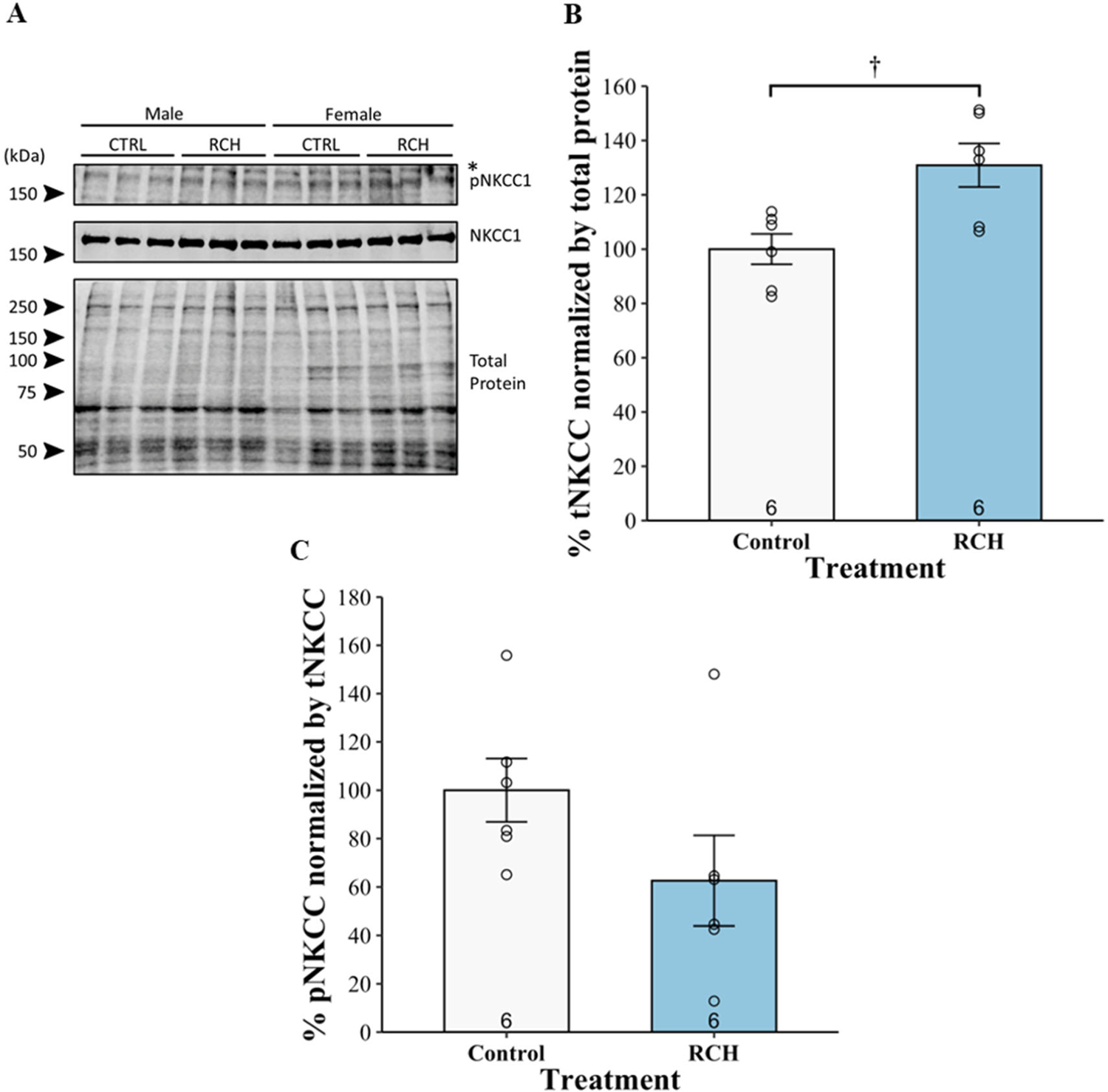
Immunoblotting of NKCC. **A**: SDS-PAGE of male and female locusts pre-treated with different conditions (control and RCH). RCH **B:** increases the protein abundance of tNKCC but **C**: does not affect the protein abundance of pNKCC. Data are displayed as means ± S.E. (n = 6 each group; p-values are reported in Results). For graphical display, data (**B** and **C**) from the two sexes are pooled as no sex differences were observed. Dagger (†) indicates a statistically significant difference between control and RCH.

## 4. Discussion

The nervous system must work properly to maintain normal body function. Extreme environmental conditions can impair neural performance (Robertson, 2004; Robertson and Money, 2012). Anoxia disrupts the energy supply, which is crucial for CNS function. The ability to tolerate anoxia is plastic and can be modulated by acute pre-treatments in many insect species. RCH is an acute pre-treatment in which brief chilling enhances insects’ ability to cope with stress. RCH delays anoxia-induced SD in locusts (Srithiphaphirom and Robertson, 2022). The present study investigates whether RCH modifies the onset of anoxia-induced SD in the locust via (1) NKA, (2) NKCC, and/or (3) K_v_ channels. The movement of ions across cell membranes is highly selective and regulated by NKA, NKCC, and K_v_ channels. As we used SD to indicate the onset of anoxia-induced neural shutdown and measured the DC potential across the BBB, we determined time to SD, recovery time from SD, and amplitude of DC potential, and further investigated NKCC phosphorylation and abundance We performed experiments on both male and female locusts in a 1:1 ratio to explore the potential effect of sex, sex-treatment interactions, and sex-drug interactions. Prior studies showed that male locusts enter coma earlier and take longer time to recover from anoxia than female locusts (Hou et al., 2014; Robertson et al., 2019; Robertson and Van Dusen, 2021; Van Dusen et al., 2020). However, in this study, there was no effect of sex, no interaction between treatment and sex, and no interaction between drug and sex in all treatments, similar to our previous study (Srithiphaphirom and Robertson, 2022).

NKA is responsible for the movement of Na^+^ and K^+^ across cell membranes against their concentration gradients via the consumption of energy. Movements of Na^+^ and K^+^ across the insect BBB involve NKA pumps to actively maintain ion homeostasis in the CNS (Treherne and Schofield, 1981). As mentioned above, the rapid surges of [K^+^]_o(CNS)_, which indicate SD, are triggered by the imbalance in processes of K^+^ accumulation and K^+^ clearance. We believe that the impairment of NKA operation may cause this imbalance. We tested whether inhibiting NKA using ouabain would affect the onset of anoxia-induced SD of locusts after RCH pre-treatment. NKA plays an important role in maintaining ion homeostasis needed for proper neuronal activity, including K^+^ clearance (Rodgers et al., 2009). We found that bath-application of ouabain decreased the time to SD of both control and RCH locusts, indicating NKA is involved in SD occurrence. This finding is well-established in mammalian models (e.g., (Balestrino et al., 1999; Menna et al., 2000; Obeidat and Andrew, 1998)). NKA is ATP-dependent, thus exposure to anoxia, which disrupts the energy supply, affects the NKA activity. The decrease in NKA activity causes a loss of ion homeostasis, which in turn triggers SD. However, reducing the time to SD of RCH locusts could not confirm that NKA takes part in the mechanism of RCH because it could be a general consequence of NKA affecting SD occurrence.

The NKCCs are a class of membrane proteins that transport Na^+^, K^+^, and Cl^-^ across the cell membrane, in most cases with a stoichiometry of 1Na^+^:1K^+^:2Cl^-^. NKCC has been shown to transport Na^+^, K^+^, and Cl^-^ into cells when [K^+^]_o_ is high, thus NKCC aids K^+^ clearance (Walz and Hertz, 1984). Blocking NKCC can cause an imbalance between K^+^ accumulation and K^+^ clearance by slowing K^+^ clearance (Spong et al., 2015). Here, we found that blocking NKCC with bumetanide only affected the time to SD of RCH locusts but not control, indicating the involvement of NKCC in the mechanism of RCH in the locust CNS.

K^+^ channels allow rapid and selective flow of K^+^ across the cell membrane, and thus, generate electrical signals in cells. One of the major classes of K^+^ channels is K_v_ channels that open or close in response to changes in the transmembrane potential. K_v_ channels play a role in the generation and propagation of electrical impulses in the nervous system and are involved in maintaining [K^+^]_o_. K_v_ channels promote K^+^ accumulation in the extracellular space (Rodgers et al., 2010). Here, we used TEA and 4-AP to block K_v_ channels. TEA is a more general K_v_ channel blocker, whereas 4-AP is a selective K_v_ blocker of the *Shaker* family. We were interested in members of the *Shaker* family because Bayley et al. (Bayley et al., 2020) found that cold acclimation (3-5 days at 11 °C) increases the transcription abundance of a *Shaker* K_v_ channel. We found that blocking K_v_ channels using TEA increased the time to SD of control locusts and had an additive effect on RCH. Our results of TEA are similar to the study by Rodgers et al. (Rodgers et al., 2009) that TEA delays the onset of ouabain-induced SD in locusts. Blocking K_v_ channels reduces K^+^ conductance, leading to a decrease in K^+^ accumulation. In contrast with Bayley et al. (Bayley et al., 2020), blocking K_v_ channels using 4-AP affected neither control nor RCH locusts. The differences between our study and Bayley et al.’s study are that (1) our pre-treatment was RCH (short-term cold hardening; 4 hours), whereas their pre-treatment was cold acclimation (long-term cold hardening; 3-5 days), and (2) we focused on ion regulation in the CNS instead of tolerance in muscle fibres. These results suggest that other families of K_v_ channels, excluding the *Shaker* family, take part in SD occurrence and the mechanism of RCH.

How does RCH regulate NKA, NKCC, and K_v_ channels? We speculated that NKA, NKCC, and K_v_ channels could be regulated by an upstream signalling pathway. Our primary suggestion is that RCH could regulate NKA, NKCC, and K_v_ channels via an octopaminergic pathway. We previously showed that octopamine (OA), an insect stress hormone, mediates the RCH-induced delay of the onset of anoxia-induced locust coma (Srithiphaphirom and Robertson, 2022). OA levels may have increased during RCH pre-treatment (mild stress) and activated a signalling pathway that includes the production of intracellular cyclic adenosine monophosphate (cAMP), protein kinase A (PKA), and protein kinase C (PKC) (Armstrong and Robertson, 2006; Evans and Robb, 1993; Robb et al., 1994; Roeder, 1992; Roeder and Gewecke, 1990). Both PKA and PKC regulate NKA and K^+^ conductance (Feschenko et al., 2000). An increase in cAMP level stimulates NKCC (Ueberschär and Bakker-Grunwald, 1985) and protein kinase can mediate protein phosphorylation of NKCC (Delpire et al., 1994; Jaggi et al., 2015). NKA also regulates the activation of NKCC. NKCC uses the electrochemical gradient established by NKA, thus a decrease in intracellular Na^+^ activates NKCC to transport Na^+^, K^+^, and Cl^-^ ions into the cell. The conformation of K_v_ channels is altered by membrane potential, triggering the opening and closing of the channels. RCH could regulate K_v_ channels by modulating K^+^ conductance via protein kinase. Finally, RCH could regulate the activity of NKA, NKCC, and K_v_ channels *via* the adjustment of membrane composition. RCH increases the amounts of oleic acid (*S. crassipalpis*: (Michaud and Denlinger, 2006)) and linoleic acid (*D. melanogaster*: (Overgaard et al., 2005)) in cell membranes which leads to an increase in membrane fluidity. This, coupled with the potential changes in ion channel localization, could affect the function of embedded NKA, NKCC, and K_v_ channels, as well as the observed abundance of NKCC in our immunoblotting experiments. However, change in membrane composition during RCH is not always detectable (MacMillan et al., 2009).

Cold acclimation causes transcriptional changes of proteins (e.g., (Bayley et al., 2020; Moribe et al., 2010; Yocum et al., 1998)). It is conceivable that RCH affected the protein expression of NKA, NKCC, and K_v_ channels although the treatment period of RCH is markedly shorter than the treatment period of cold acclimation. Because the pharmacological experiments clearly indicated a role for NKCC in RCH, we performed immunoblotting against total NKCC (tNKCC) and phosphorylated NKCC (pNKCC) with and without prior RCH. We found that RCH increased the abundance of NKCC but did not significantly affect the phosphorylation level. We note that pNKCC tended to decrease after RCH and the lack of statistical significance appears to be driven by the result for one sample, so this negative result should be interpreted cautiously. Nevertheless, the combination of increased tNKCC with no change in phosphorylation would be expected to decrease pNKCC abundance relative to tNKCC. The increased NKCC abundance after RCH pre-treatment would help maintain [K^+^]_o(CNS)_ during anoxia-induced coma and prevent cell swelling, which is associated with increased severity of SD indicated by more rapid onset and larger amplitude events (Spong et al., 2015).

The mechanism underlying the increase in NKCC abundance remains unclear. It is generally accepted that the effects of RCH without a recovery period are most likely not due to increased transcription because of the short time course and low temperatures of exposure (Sinclair et al., 2007; Teets et al., 2020). In contrast to those studies, we did not perform the experiments immediately after RCH pre-treatment and thus it is possible that the protein expression of NKCC increased during the hour at RT before the electrophysiology experiments. However, RCH without a recovery period does delay anoxic coma and ouabain-induced SD in locusts (Gantz et al., 2020) suggesting that the electrophysiological results of our experiments may not have been caused by increased protein expression. RCH may have altered NKCC protein turnover reducing the rate of NKCC degradation and leading to increased abundance. On balance, we currently favour the interpretation that RCH increased the abundance of NKCC by reducing its degradation.

Considering the recovery time from anoxia-induced SD and the amplitude of DC potential, ouabain had no effect on the amplitude of DC potential but increased the recovery time from SD of control locusts. Recovery from SD involves the re-establishment of ion homeostasis in the nervous system (Robertson et al., 2017). NKA aids K^+^ clearance, thus inhibiting NKA would hinder the re-establishment of ion homeostasis, and consequently slow down recovery from SD. However, ouabain did not significantly increase the recovery time from SD of RCH locusts. Our results are similar to the study of heat shock (HS) pre-treatment by Rodgers et al. (Rodgers et al., 2007). Ouabain increases time to recovery in control locusts but not in HS locusts. Here, based on the results of control and RCH locusts with saline, RCH did not have a direct effect on the recovery time from SD. However, NKA could be more resilient after RCH pre-treatment thus becoming less sensitive to ouabain. TEA had no effect on the recovery time from SD but decreased the amplitude of DC potential of both control and RCH locusts. Prior studies showed that TEA reduces the amplitude of the [K^+^]_o(CNS)_ in locusts (Rodgers et al., 2007; Rodgers et al., 2009) and hypoxic SD amplitude of rat hippocampal slices (Aitken et al., 1991). It is possible that blocking K_v_ channels, in general, inhibits K^+^ movement and slows K^+^ accumulation thus affecting the magnitude of ion disturbance. It should be noted that we measured the amplitude of DC potential at a single point where we inserted the microelectrode, so it may not accurately represent the overall ion disturbance. Bumetanide and 4-AP affected neither recovery time from SD nor amplitude of DC potential.

In summary, using a pharmacological approach, our results demonstrate that in the locust CNS (1) NKA is involved in SD occurrence and may take part in the mechanism of RCH, (2) NKCC is involved in the mechanism of RCH, and (3) K_v_ channels, excluding the *Shaker* family, take part in SD occurrence and the mechanism of RCH. We propose that RCH could regulate NKA, NKCC, and K_v_ channels through the octopaminergic pathway (Srithiphaphirom and Robertson, 2022). This study sheds light on the mechanism of RCH and its effect on the regulation of K^+^ homeostasis in the CNS during anoxia. Neither NKA nor K_v_ channels play important roles in RCH. However, the mechanism of RCH likely involves regulating the abundance of NKCC but not its phosphorylation state. Thus, future studies could validate and confirm the mechanism of changes in NKCC level following RCH and characterize the involvement and regulation of other ion channels and transporters in this response.

### Grants

This study was supported by a Discovery grant to RMR (#04561-2017) from the Natural Sciences and Engineering Research Council of Canada (NSERC).

## Disclosures

The authors declare no conflicts of interest, financial or otherwise.

## Author Contributions

Phinyaphat Srithiphaphirom: Methodology, Investigation, Formal analysis, Visualization, Writing – original draft, Writing – review & editing. Yuyang Wang: Methodology, Investigation, Writing – review & editing. Maria J. Aristizabal: Resources, Supervision, Writing – review and editing. R. Meldrum Robertson: Conceptualization, Resources, Funding acquisition, Supervision, Writing – review & editing.

## Notes

### Competing Interest Statement

The authors have declared no competing interest.

