## Supplementary Data for "Rapid cold hardening modifies mechanisms of ion regulation to delay anoxia-induced spreading depolarization in the CNS of *Locusta migratoria*"

**Table S1. RCH and pharmacological agent experiments.** Time to anoxia-induced SD, recovery time from anoxia-induced SD, and amplitude of DC potential in semi-intact preparation. Parametric data are presented as Mean ± Standard Error; nonparametric data are presented as Median (Inter Quartile Range). Data from the two sexes are pooled as no sex differences were observed.

| | | n | Time to SD (min) | Recovery time (min) | Amplitude of DC (mV) |
|---|---|---|---|---|---|
| Control | Saline | 10 | 5.0 (3.7 – 5.3) | 1.6 ± 0.1 | 55.1 ± 1.6 |
| | Ouabain | 10 | 1.5 (1.1 – 1.8) | 2.0 ± 0.2 | 57.1 ± 0.9 |
| RCH | Saline | 10 | 7.8 (6.2 – 9.2) | 1.5 ± 0.1 | 56.9 ± 1.4 |
| | Ouabain | 10 | 2.5 (1.8 – 3.8) | 1.8 ± 0.2 | 56.9 ± 1.2 |
| Control | Saline | 10 | 4.3 ± 0.4 | 1.6 ± 0.1 | 54.7 ± 1.4 |
| | Bumetanide | 10 | 3.9 ± 0.3 | 1.4 ± 0.1 | 54.0 ± 1.4 |
| RCH | Saline | 10 | 8.0 ± 0.6 | 1.5 ± 0.1 | 57.5 ± 1.4 |
| | Bumetanide | 10 | 5.9 ± 0.3 | 1.5 ± 0.1 | 52.0 ± 1.6 |
| Control | Saline | 10 | 4.5 ± 0.3 | 1.5 ± 0.1 | 54.0 ± 1.5 |
| | TEA | 10 | 6.1 ± 0.3 | 1.5 ± 0.1 | 43.6 ± 1.3 |
| RCH | Saline | 10 | 7.2 ± 0.3 | 1.4 ± 0.1 | 55.8 ± 1.2 |
| | TEA | 10 | 10.4 ± 0.7 | 1.4 ± 0.1 | 46.5 ± 1.4 |
| Control | Saline | 10 | 4.5 ± 0.3 | 1.5 ± 0.1 | 54.0 ± 1.5 |
| | 4-AP | 10 | 4.4 ± 0.3 | 1.5 ± 0.1 | 57.3 ± 1.3 |
| RCH | Saline | 10 | 7.2 ± 0.3 | 1.4 ± 0.1 | 55.8 ± 1.2 |
| | 4-AP | 10 | 6.3 ± 0.3 | 1.4 ± 0.1 | 57.3 ± 1.8 |

**Table S2. Percentage of total NKCC (tNKCC) normalized by total protein and percentage of phosphorylated NKCC (pNKCC) normalized by tNKCC.** Data are presented as Mean ± Standard Error. Data from the two sexes are pooled as no sex differences were observed.

| | n | Total NKCC (%) | Phospho-NKCC (%) |
|---|---|---|---|
| Control | 6 | 100.0 ± 5.6 | 100.0 ± 13.1 |
| RCH | 6 | 130.9 ± 8.0 | 62.6 ± 18.7 |

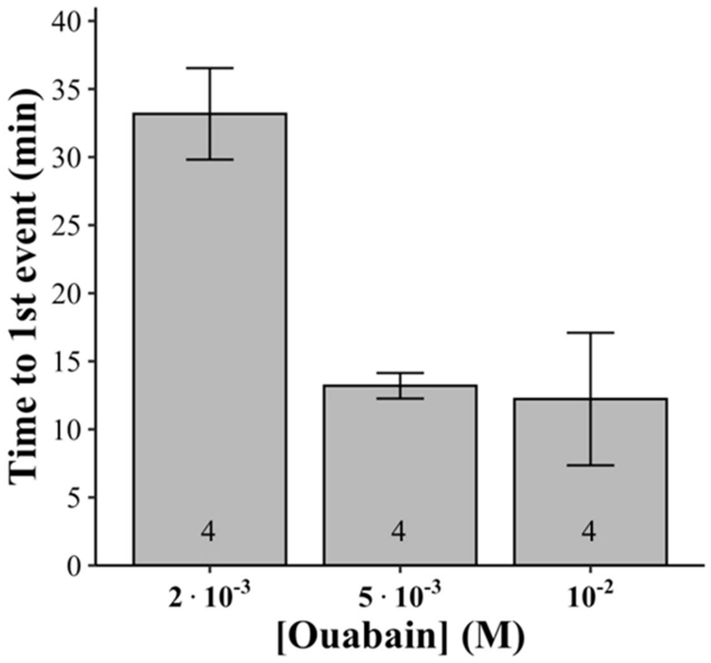


**Figure S1. Dose-response curve of ouabain.** No anoxia was required. $10^{-2}$ M ouabain, as previously used (Van Dusen et al., 2020), gave times to the first SD event of three individuals of 7.2, 5.9, and 9.1 minutes with one individual that had the first event after 25 minutes. $5 \times 10^{-3}$ M ouabain gave times to the first SD event of 15.1, 12.5, 14.3, and 10.9 minutes. $2 \times 10^{-3}$ M ouabain gave times to the first SD event of 39.2, 25.7, 38.5, 29.3 minutes. Based on these results, we decided to use $5 \times 10^{-3}$ M ouabain with a treatment duration of 10 minutes. In this figure, data are displayed as means ± S.E. (n = 4 each group). For graphical display, data from the two sexes are pooled as no sex differences were observed.

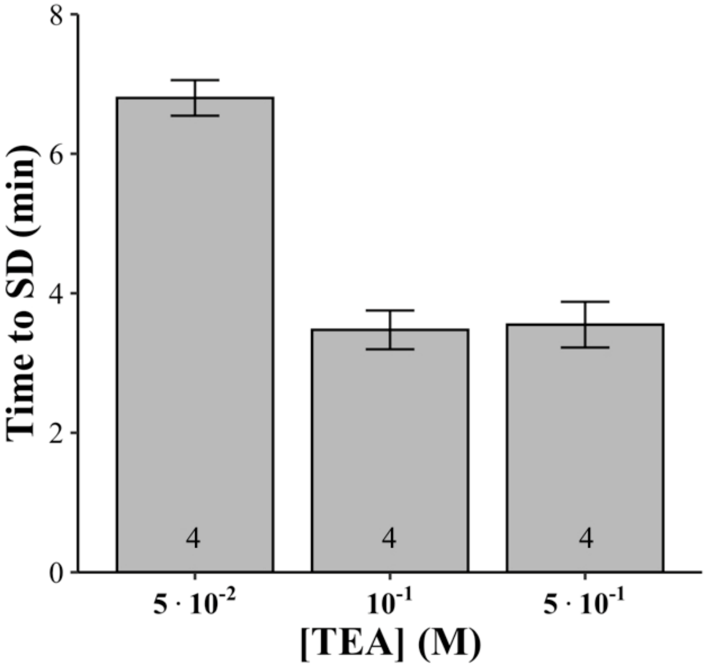


**Figure S2. Dose-response curve of TEA.** We used anoxia to induce SD. $10^{-1}$ M TEA, as previously used (Rodgers et al., 2007), gave times to the first SD event of four individuals of 4.0, 3.5, 2.7, 3.7 minutes. $5x10^{-1}$ M TEA gave time to first SD event of 3.4, 3.3, 3.0, 4.5 minutes. $5x10^{-2}$ M TEA gave times to first SD event of 6.3, 7.5, 6.8, 6.6 minutes. Based on these results, we decided to use $5x10^{-2}$ M TEA with a treatment duration of 15 minutes (Rodgers et al., 2007). In this figure, data are displayed as means ± S.E. (n = 4 each group). For graphical display, data from the two sexes are pooled as no sex differences were observed.

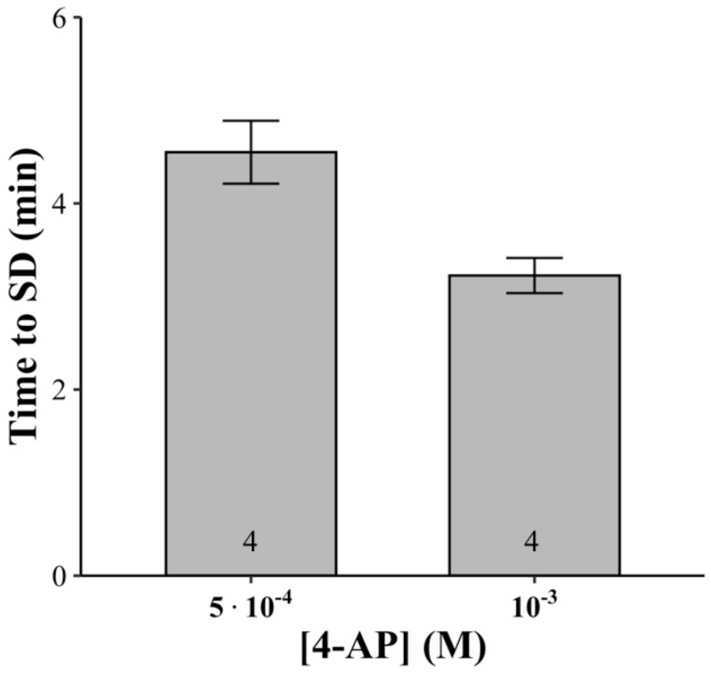


**Figure S3. Dose-response curve of 4-AP.** We used anoxia to induce SD. $10^{-3}$ M 4-AP, as previously used (Bayley et al., 2020), gave times to the first SD event of four individuals of 3.6, 2.7, 3.3, and 3.3 minutes. $5x10^{-4}$ M 4-AP gave times to SD of four individuals of 5.4, 3.8, 4.7, and 4.3 minutes. Based on these results, we decided to use $5x10^{-4}$ M 4-AP. We used the same treatment duration as the TEA experiment (15 minutes). Data are displayed as means ± S.E. (n = 4 each group). For graphical display, data from the two sexes are pooled as no sex differences were observed.

Bayley, J.S., Sørensen, J.G., Moos, M., Koštál, V., Overgaard, J., 2020. Cold acclimation increases depolarization resistance and tolerance in muscle fibers from a chill-susceptible insect, *Locusta migratoria*. Am J Physiol Regul Integr Comp Physiol 319, R439-r447.

Rodgers, C.I., Armstrong, G.A.B., Shoemaker, K.L., LaBrie, J.D., Moyes, C.D., Robertson, R.M., 2007. Stress preconditioning of spreading depression in the locust CNS. Plos One 2, e1366.

Van Dusen, R.A., Shuster-Hyman, H., Robertson, R.M., 2020. Inhibition of ATP-sensitive potassium channels exacerbates anoxic coma in *Locusta migratoria*. J Neurophysiol 124, 1754-1765.
